# Inhibition of protein tyrosine phosphatase PTP1B function ameliorates pathophysiological deficits in Rett Syndrome

**DOI:** 10.64898/2026.06.04.730096

**Authors:** Christopher A. Bonham, Christy Felice, Lisa N. Christensen, Nicholas K. Tonks

**Affiliations:** Cold Spring Harbor Laboratory; Cold Spring Harbor, NY 11724, USA

**Author notes:** Department of Biochemistry, Schulich School of Medicine & Dentistry, Western University; London, ON N6A 3K7, Canada.

## Abstract

Rett syndrome (RTT) is a severe neurodevelopmental disorder in which current therapeutic strategies remain largely focused on providing symptomatic relief without addressing underlying disease mechanisms. In contrast, we have identified the protein tyrosine phosphatase PTP1B as a mechanism-based therapeutic target and evaluated a class of selective, allosteric small-molecule inhibitors in female murine models of RTT. We show that one of these compounds localizes to brain regions central to motor coordination and cardio-respiratory control, which are core domains of RTT pathology. Pharmacological inhibition of PTP1B produces robust and sustained improvement in multiple disease symptoms, including muscle weakness, motor and coordination deficits, and cardiac and respiratory dysfunction. Concordant results obtained with genetic ablation of PTP1B, with effects maintained for over one year, demonstrate that phenotypic rescue arises from on-target modulation of disease-relevant signaling. Mechanistically, PTP1B inhibition is known to normalize neurotrophic and metabolic pathways, including TRKB and insulin/leptin signaling, thereby restoring circuit-level function. These findings establish PTP1B as a clinically actionable, disease-modifying target and demonstrate that selective, allosteric inhibition of a protein tyrosine phosphatase can achieve durable therapeutic benefit in vivo. This work provides a strong rationale for the clinical evaluation of PTP1B inhibitors as a mechanism-based treatment strategy for RTT.

**One Sentence Summary:** We have validated inhibition of PTP1B as a mechanism-based therapeutic strategy that alleviates a wide range of symptoms in a mouse model of Rett syndrome.

## INTRODUCTION

Rett syndrome (RTT) is a progressive X-linked neurodevelopmental disorder that primarily affects females and is most commonly caused by mutations in the transcriptional regulator methyl-CpG binding protein 2 (MECP2). Affected individuals exhibit developmental regression beginning around 6-18 months of age, with features including loss of motor and communication skills, repetitive hand movements, impaired coordination, microcephaly, and severe complications such as cardiac arrhythmias, respiratory dysfunction, and seizures [1–5]. Patients typically survive into mid-adulthood but require lifelong care, and effective disease-modifying therapies remain lacking.

Loss of MECP2 function at any stage of life produces RTT-like phenotypes, indicating a continuous requirement for MECP2 in maintaining normal neuronal function [6–8]. Notably, restoration of MECP2 expression in adult mouse models reverses key disease features [9–10], demonstrating that RTT phenotypes are not irrevocable and may be therapeutically modified. However, the mosaic expression of MECP2 and the narrow tolerance for its dosage, where both deficiency and excess are pathogenic, pose substantial challenges for gene replacement strategies [11–12]. These limitations underscore the need for alternative therapeutic approaches, particularly those based on small molecules capable of modulating downstream disease mechanisms [13–14].

Disruption of neurotrophic signaling has emerged as a central feature of RTT. Decreased signaling through the receptor tyrosine kinase TRKB, the receptor for brain-derived neurotrophic factor (BDNF), contributes to neuronal dysfunction, and enhancing TRKB activity has been associated with phenotypic improvement [15]. Although direct TRKB agonists have been explored [6], targeting negative regulators of TRKB signaling may provide a more physiologically responsive strategy to rescue signaling. Previously, we identified *PTPN1*, the gene encoding the protein tyrosine phosphatase PTP1B, as a direct transcriptional target of MECP2, with loss of MECP2 function leading to increased PTP1B expression [16]. Elevated PTP1B suppresses TRKB phosphorylation and impairs BDNF signaling, whereas inhibition of PTP1B restores TRKB activation and downstream signaling in RTT models, supporting PTP1B as a mechanism-based therapeutic target [16].

Here, we have developed a preclinical platform to evaluate the longitudinal effects of selective, allosteric small-molecule inhibitors of PTP1B in female mouse models of RTT. These compounds localize to cortical and brain stem regions governing motor and cardio-respiratory function, key domains of RTT pathology. We demonstrate that both pharmacological inhibition and genetic ablation of PTP1B produced robust and sustained improvements in motor function, brain weight, and cardio-respiratory physiology. Convergent genetic and pharmacological evidence indicates that these effects arose from on-target modulation of disease-relevant signaling pathways. Together, these findings establish PTP1B as a clinically actionable, disease-modifying target and support the therapeutic potential of selectively modulating phosphatase-dependent signaling in RTT. More broadly, these findings define a mechanistic framework in which dysregulated phosphatase activity contributes to neurological disease and can be therapeutically corrected. This work therefore supports the broader concept that selective targeting of protein tyrosine phosphatases may represent a generalizable strategy for modulating signal transduction in disorders of the central nervous system.

## RESULTS

### Optimization of phenotypic analysis of PTP1B inhibition in RTT mouse models

Genetic mouse models of Rett syndrome (RTT) that recapitulate key behavioral features of the disease have enabled preclinical therapeutic evaluation [17–20]. Restoration of Mecp2 expression has been shown to reverse RTT-like phenotypes [9–10], indicating that these models provide a tractable platform for testing disease-modifying interventions.

We established a robust, longitudinal platform to evaluate the impact of PTP1B inhibition in RTT mouse models, incorporating standardized cohort stratification, expanded phenotypic assays, and defined assessment timelines (Fig. 1A). Female animals were assigned to experimental groups based on genotype, body weight, behavioral performance, and litter, yielding highly consistent baseline characteristics across independent cohorts (Fig. 1, B & C; fig. S1, A-D). Pre-study acclimatization minimized variability associated with routine handling, restraint, compound administration, and experimental apparatus.

**Figure 1.**
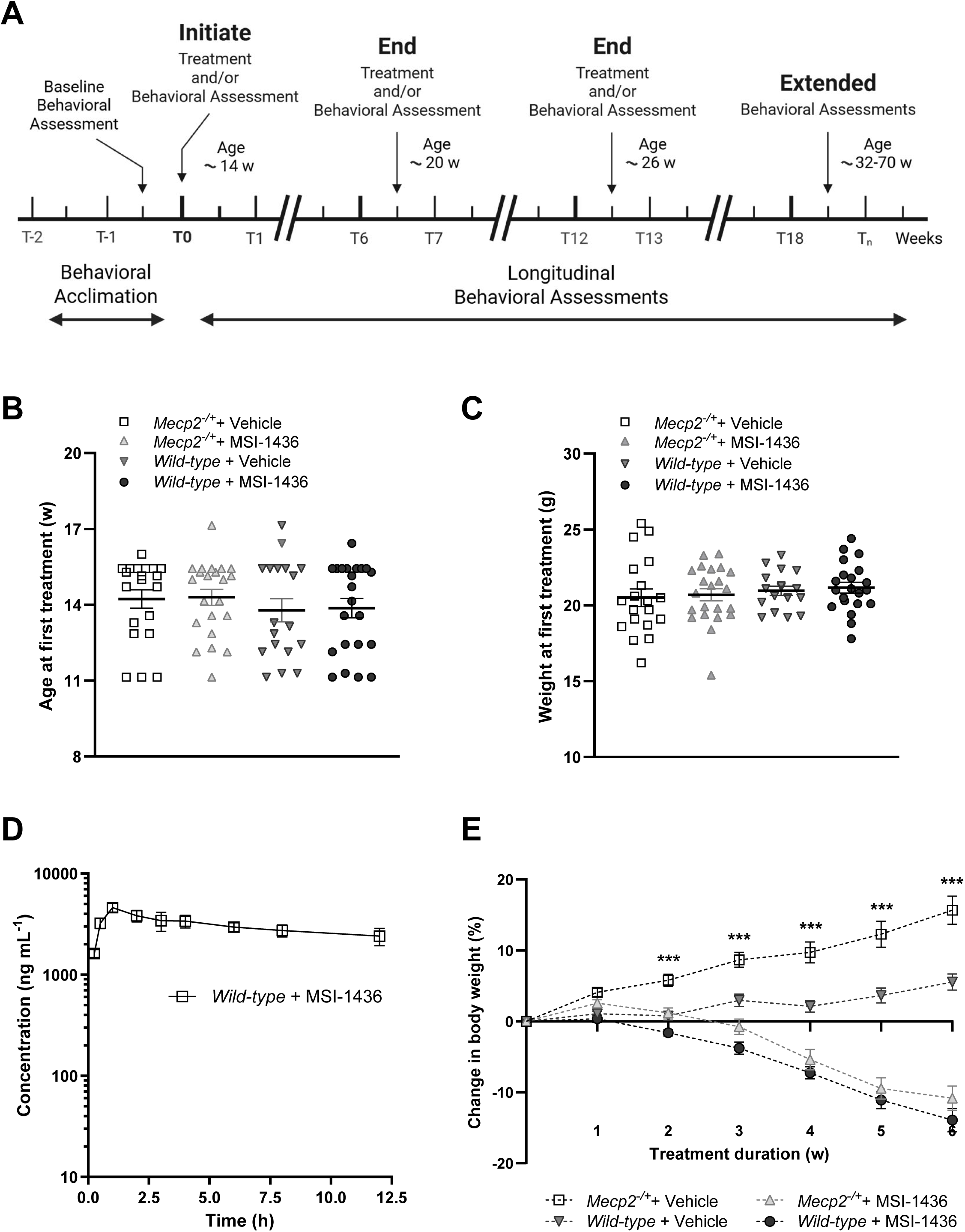
A longitudinal platform for assessing on-target inhibition of PTP1B in a murine model of Rett Syndrome. (**A**) Schematic overview of the experimental design. Female mice (12-14 weeks of age) underwent a 2-week acclimatization period, followed by baseline physical, behavioral, and physiological assessments to establish balanced cohorts. Longitudinal treatment (intraperitoneal injection with MSI-1436 every 72 hours at 1-10 mg/kg dose) and serial phenotyping were then performed over 2-12 months. Schematic created with BioRender.com. (**B**) Age distribution at first treatment across cohorts (*Mecp2^−/+^* + vehicle, *n* = 20; *Mecp2^−/+^*+ MSI-1436, *n* = 22; wild-type + vehicle, *n* = 18; wild-type + MSI-1436, *n* = 22). (**C**) Weight distribution at first treatment across cohorts (*Mecp2^−/+^* + vehicle, *n* = 19; *Mecp2^−/+^* + MSI-1436, *n* = 22; wild-type + vehicle, *n* = 17; wild-type + MSI-1436, *n* = 21). Mice lacking equivalent pre-treatment body weight measurements were excluded from analysis (*Mecp2^−/+^*+ Vehicle, *n* = 1; wild-type + Vehicle, *n* = 1; wild-type + MSI-1436, *n* = 1). (**D**) Pharmacokinetics of MSI-1436 following a single intraperitoneal injection (2 mg/kg), shown as the mean blood plasma concentration at each time point (wild-type + MSI-1436, *n* = 3). One animal with concentrations > 50-fold lower than the mean of the remaining animals across multiple time points was excluded from analysis (*n* = 1). (**E**) Longitudinal change in body weight, expressed as the percent change in mean body weight per animal per week (*Mecp2^−/+^* + vehicle, *n* = 19; *Mecp2^−/+^* + MSI-1436, *n* = 22; wild-type + vehicle, *n* = 17; wild-type + MSI-1436, *n* = 21). Significance denotes comparison between both *Mecp2^−/+^* + vehicle and *Mecp2^−/+^* + MSI-1436, and *Mecp2^−/+^*+ vehicle and wild-type + vehicle. Mice lacking equivalent pretreatment body weight measurements were excluded from analysis as in (C), (*Mecp2^−/+^* + Vehicle, *n* = 1; wild-type + Vehicle, *n* = 1; wild-type + MSI-1436, *n* = 1). Data are presented as the mean ± SEM. Statistical significance was determined using unpaired *t*-tests * *p*< 0.05, ** *p*< 0.01, *** *p*< 0.001.

Study design was informed by pharmacokinetic and pharmacodynamic parameters. Following administration of the PTP1B inhibitor MSI-1436, compound half-life (t1/2 15–20 h) and time to peak concentration (Tmax 0.5–2 h) supported an intraperitoneal (i.p.), twice-weekly regimen of 2 mg kg⁻¹ dosing, with phenotypic assessments performed ∼24 h post-dose (Fig. 1D and fig. S1E).

Considering the established role of PTP1B in insulin and leptin signaling [21], we assessed metabolic phenotypes in *Mecp2^⁻/+^* mice. These animals exhibited age-dependent weight gain relative to wild-type littermates, consistent with impaired metabolic regulation, and with insulin resistance also noted in some RTT patients [22–24]. Genetic studies have demonstrated that PTP1B-null mice are healthy, display enhanced insulin sensitivity, do not develop type 2 diabetes, and are resistant to obesity when fed with a high fat diet [21]. We observed that treatment with MSI-1436 prevented weight gain in Mecp2⁻/+ mice (Fig. 1E), consistent with on-target inhibition of PTP1B and improved metabolic homeostasis. Together, these data validate a robust and reproducible platform for longitudinal evaluation of PTP1B-targeted interventions in RTT.

### Pharmacological inhibition of PTP1B improves motor and behavioral phenotypes

Although studies using hemizygous male *Mecp2^−/y^* mice avoid problems of mosaicism of X chromosomal inactivation, this complete deficiency of MECP2 does not reflect the natural history of the disease in human patients because of their short lifespan and phenotype severity. Female Mecp2^⁻/+^ mice provide a physiologically relevant model of RTT due to mosaic MECP2 expression and disease progression that more closely reflects human pathology [25]. Building on prior evidence that PTP1B is transcriptionally regulated by MECP2 and suppresses TRKB signaling [16], we evaluated the effects of sustained PTP1B inhibition in these animals across a battery of behavioral and physiological metrics (Fig. 2A).

**Figure 2.**
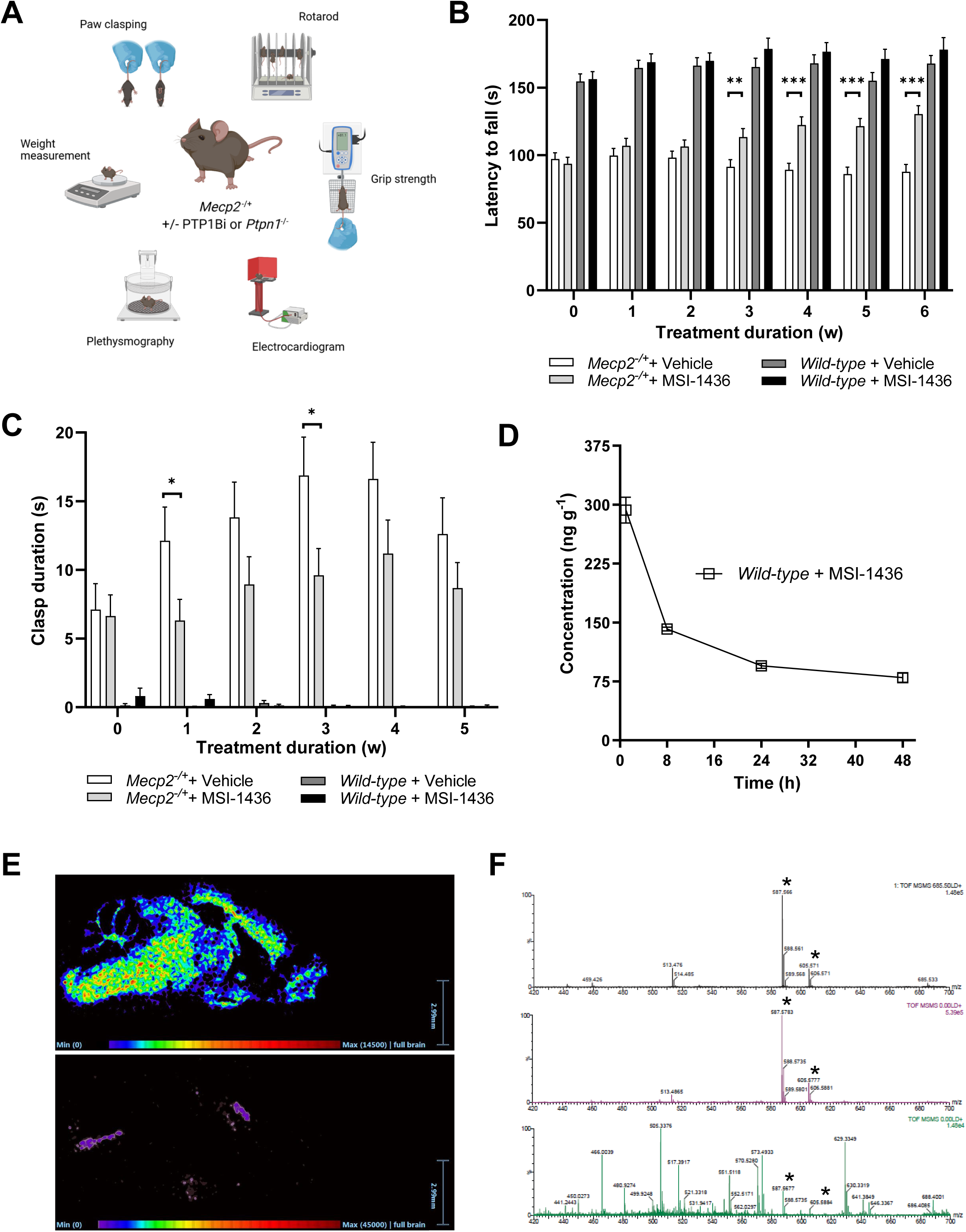
The PTP1B inhibitor MSI-1436 penetrates the brain and improves motor and coordination deficits in *Mecp2^−/+^* mice. (**A**) A schematic illustration of the longitudinal phenotyping pipeline used to monitor physical, behavioral, and physiological outcomes across treatment cohorts. Schematic created with BioRender.com. (**B**) Rotarod performance, measured as latency to fall and expressed as the mean of four trials per animal per week (*Mecp2^−/+^* + vehicle, *n* = 20; *Mecp2^−/+^* + MSI-1436, *n* = 22; wild-type + vehicle, *n* = 18; wild-type + MSI-1436, *n* = 22; 0-300 s, all trials). Mice lacking equivalent measurements were excluded from analysis for the indicated time point (*Mecp2^−/+^* + vehicle, *n* = 1, Week 6; *Mecp2^−/+^* + MSI-1436, *n* = 1, Week 6). (**C**) Hind-limb clasping, measured as mean clasp duration from two trials per animal per week (*Mecp2^−/+^*+ vehicle, *n* = 20; *Mecp2^−/+^* + MSI-1436, *n* = 22; wild-type + vehicle, *n* = 18; wild-type + MSI-1436, *n* = 22; all trials were of 45 seconds duration). Mice lacking equivalent measurements were excluded from analysis for the indicated time point (*Mecp2^−/+^* + vehicle, *n* = 1, Week 0; wild-type + Vehicle, *n* = 12, Week 0; wild-type + MSI-1436, *n* = 14, Week 0). In B & C, evaluations were initiated at a mean age of 14 weeks. (**D**) Brain pharmacokinetics of MSI-1436 following a single intravenous injection per animal (2 mg/kg) shown as the mean brain concentration at each time point (wild-type + MSI-1436, *n* = 24; *n* = 6 per time point). (**E**) Representative mass spectrometry image showing the spatial distribution of a neuronal marker (*m/z* 866.6; top) and MSI-1436 (*m/z* 685.5; bottom) in brain sections of *Mecp2^−/+^*mice collected 2-3 hours after the final intraperitoneal 10 mg/kg dose. The following samples were analyzed: *Mecp2^−/+^*+ vehicle, *n* = 2; *Mecp2^−/+^*+ MSI-1436, *n* = 2; wild-type + vehicle, *n* = 2; wild-type + MSI-1436, *n* = 2). (**F**) Representative tandem mass spectrometry imaging of purified MSI-1436 (*m/z* 685.5) spotted onto a MALDI target plate (top), on adjacent tissue (middle), and of compound distribution in the brain of *Mecp2^−/+^*mice (bottom), collected 2-3 hours following the final intraperitoneal 10 mg/kg dose. (*Mecp2^−/+^* + vehicle, *n* = 2; *Mecp2^−/+^* + MSI-1436, *n* = 2; wild-type + Vehicle, *n* = 2; wild-type + MSI-1436, *n* = 2). Data represent the mean ± SEM. Statistical significance was determined using unpaired *t*-tests * *p*< 0.05, ** *p*< 0.01, *** *p*< 0.001.

Treatment with MSI-1436 resulted in progressive improvement in motor coordination, as assessed by rotarod performance, and reduced hind-limb clasping across three independent cohorts (Fig. 2, B & C; fig. S1). These effects were sustained over a 6-week treatment period and were not observed in wild-type mice, indicating disease-specific efficacy.

Pharmacokinetic analysis confirmed durable compound exposure in the brain, with detectable levels persisting for 24–48 h post-treatment (Fig. 2D). Spatial mapping demonstrated localization to the medial prefrontal cortex and medulla, regions critical for motor and cardio-respiratory control (Fig. 2, E & F) [26]. Together, these findings demonstrate that pharmacological inhibition of PTP1B produces sustained, regionally targeted, and disease-specific improvements in RTT-relevant phenotypes.

### Genetic ablation of PTP1B confirms on-target therapeutic effects

To determine whether these effects reflect on-target modulation of PTP1B, we performed genetic studies by crossing Mecp2^⁻/+^ mice with *Ptpn*1-knockout animals (Fig. 3A). As expected, *Mecp2^−/+^* mice progressively gained weight, resulting in a 15% increase in body weight over this 6-week time-period, consistent with previous observations (Fig. 3 B & C and Fig. 1E). Deletion of PTP1B in Mecp2^⁻/+^ mice significantly reduced pathological weight gain (>30%) without affecting wild-type controls (Fig. 3C). Motor performance was similarly improved, with PTP1B-deficient Mecp2⁻/+ mice showing sustained rotarod performance over 6 weeks compared to progressive decline in Mecp2⁻/+ mice (Fig. 3D). Improvements in hind-limb clasping (>65%) and grip strength (∼15%) further supported functional rescue (Fig. 3, E & F).

**Figure 3.**
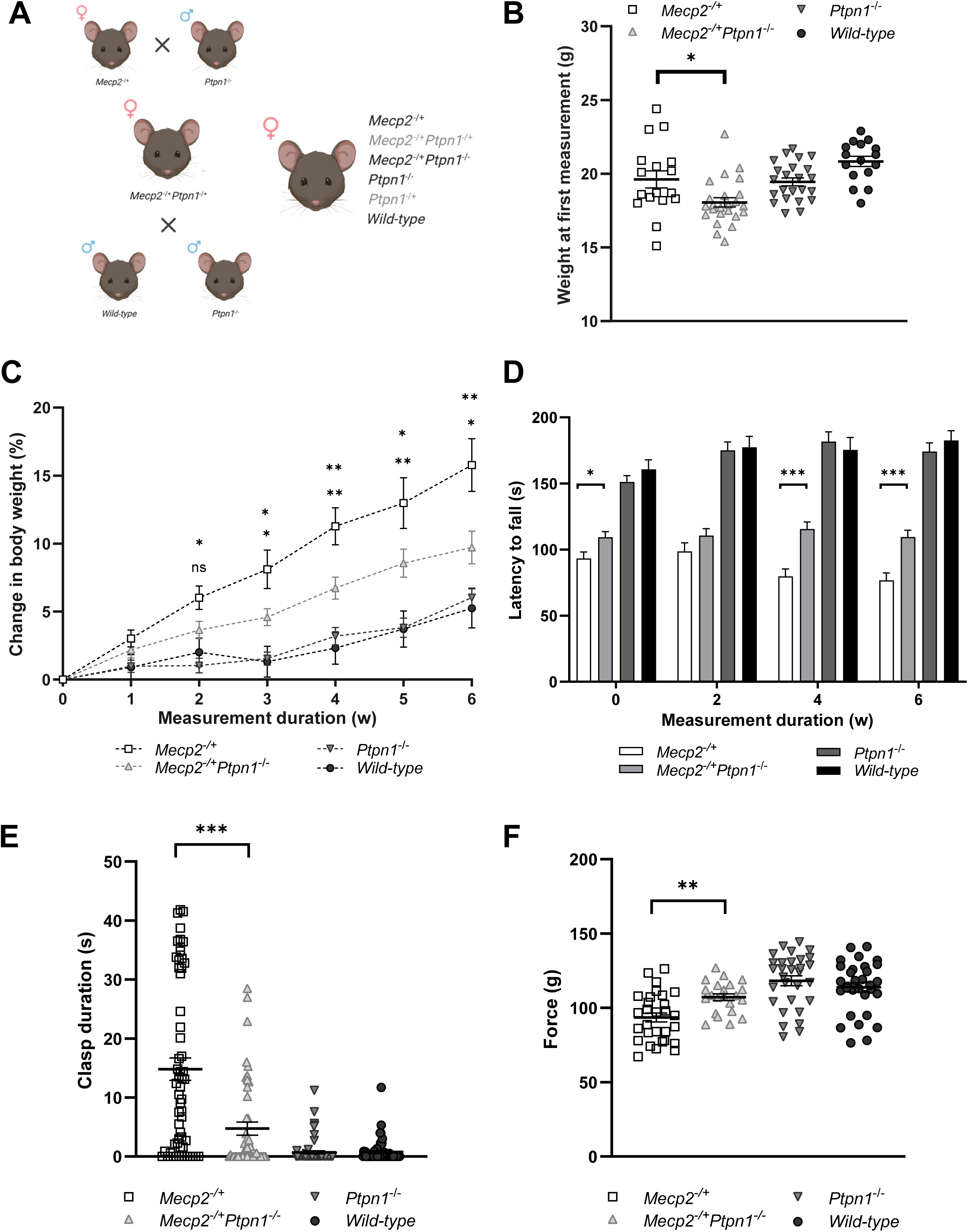
Genetic deletion of PTP1B phenocopies behavioral improvements in *Mecp2^−/+^* mice treated with PTP1B inhibitor. (**A**) Breeding strategy used to generate female mice of the indicated genotypes. Female *Mecp2^−/+^* mice were crossed with male *Ptpn1^−/-^* mice to generate female *Mecp2^−/+^Ptpn1^−/+^* offspring, which were subsequently bred with male *Ptpn1^−/-^*and wild-type mice to generate the following genotypes: *Mecp2^−/+^*, *Mecp2^−/+^Ptpn1^−/+^*, *Mecp2^−/+^Ptpn1^−/-^*, *Ptpn1^−/+^*, *Ptpn1^−/-^*, and wild-type female mice. (**B**) Body weight distribution at first measurement. Evaluations were initiated at a mean age of 12-13 weeks (B – F). (**C**) Longitudinal change in body weight expressed as the percent change in body weight per animal per week (*Mecp2^−/+^*, *n* = 17; *Mecp2^−/+^Ptpn1^−/-^*, *n* = 24; *Ptpn1^−/-^*, *n* = 23; wild-type, *n* = 16). Significance denotes comparison between *Mecp2^−/+^* and *Mecp2^−/+^Ptpn1^−/-^* (top), and between wild-type and *Mecp2^−/+^Ptpn1^−/-^* (bottom). Mice lacking equivalent body weight measurements were excluded from analysis for the indicated time point (*Mecp2^−/+^*, *n* = 3, Week 3; *Mecp2^−/+^Ptpn1^−/-^*, *n* = 5, Week 3; *Ptpn1^−/-^*, *n* = 4, Week 3; wild-type, *n* = 2, Week 3). One *Mecp2^−/+^* mouse with recurrent body weight fluctuations attributed to malocclusion was excluded from analysis. (**D**) Rotarod performance, measured as latency to fall and expressed as the mean of four trials per animal per week (*Mecp2^−/+^*, *n* = 17; *Mecp2^−/+^Ptpn1^−/-^*, *n* = 24; *Ptpn1^−/-^*, *n* = 23; wild-type, *n* = 16). Range 0-300 seconds for all trials. One *Mecp2^−/+^* mouse with measurements repeatedly identified as an outlier over multiple weeks was excluded from analysis. (**E**) Hind-limb clasping at first measurement, expressed as the mean clasp duration from two trials per animal (*Mecp2^−/+^*, *n* = 29; *Mecp2^−/+^Ptpn1^−/-^*, *n* = 23; *Ptpn1^−/-^*, *n* = 31; wild-type, *n* = 29). Range 0-45 seconds for all trials. (**F**) Forelimb grip strength at the first measurement, expressed as the mean of five trials per animal (*Mecp2^−/+^*, *n* = 29; *Mecp2^−/+^Ptpn1^−/-^*, *n* = 22; *Ptpn1^−/-^*, *n* = 30; wild-type, *n* = 31). Data represent the mean ± SEM. Outliers were evaluated by the Rout method (Q = 1%). Statistical significance was determined using unpaired *t*-tests * *p*< 0.05, ** *p*< 0.01, *** *p*< 0.001.

A gene-dosage effect was observed, as heterozygous deletion of PTP1B partially attenuated phenotypes (fig. S2 A-D), indicating that precise modulation of PTP1B activity is required for optimal therapeutic benefit. Importantly, PTP1B deletion had no effect in wild-type animals (Fig. 3D), demonstrating that observed improvements are specific to correction of RTT-associated deficits. Concordance between pharmacological inhibition and genetic ablation provides strong evidence that phenotypic rescue arises from on-target modulation of PTP1B-dependent signaling.

### PTP1B ablation produces durable improvements in neurodevelopment and function

Post-mortem studies have revealed that RTT patients display a decrease in brain weight and volume (microcephaly), up to 35% lower than normal matched controls [27–29]. Similar results are reported from animal models [30,31]. This reduced brain size is due to impaired neurodevelopment rather than neurodegeneration, raising the possibility that these effects may be reversible. PTP1B deletion in *Mecp2^⁻/+^* mice significantly increased brain weight at 24 and 45 weeks of age compared to *Mecp2^⁻/+^* controls, while limiting pathological body weight gain (Fig. 4, A–D). The average difference in brain weight more than doubled (24 mg) by age ∼45 weeks, whereas the brain weight difference between *Ptpn1^−/-^* and *wild-type* mice was < 5 mg on average across these same age ranges (Fig. 4, B-D). At ∼65 weeks of age, no difference in brain weight between *Mecp2^−/+^* and *Mecp2^−/+^Ptpn1^−/-^*mice is observed; however, at this age, *Mecp2^−/+^* mice exhibit an extreme obesity phenotype, which may instead contribute to masking of a brain weight phenotype rather than represent neurodevelopmental recoveries (fig. S2, E & F). These effects were not observed in Ptpn1^⁻/⁻^ mice alone, indicating disease-specific rescue.

**Figure 4.**
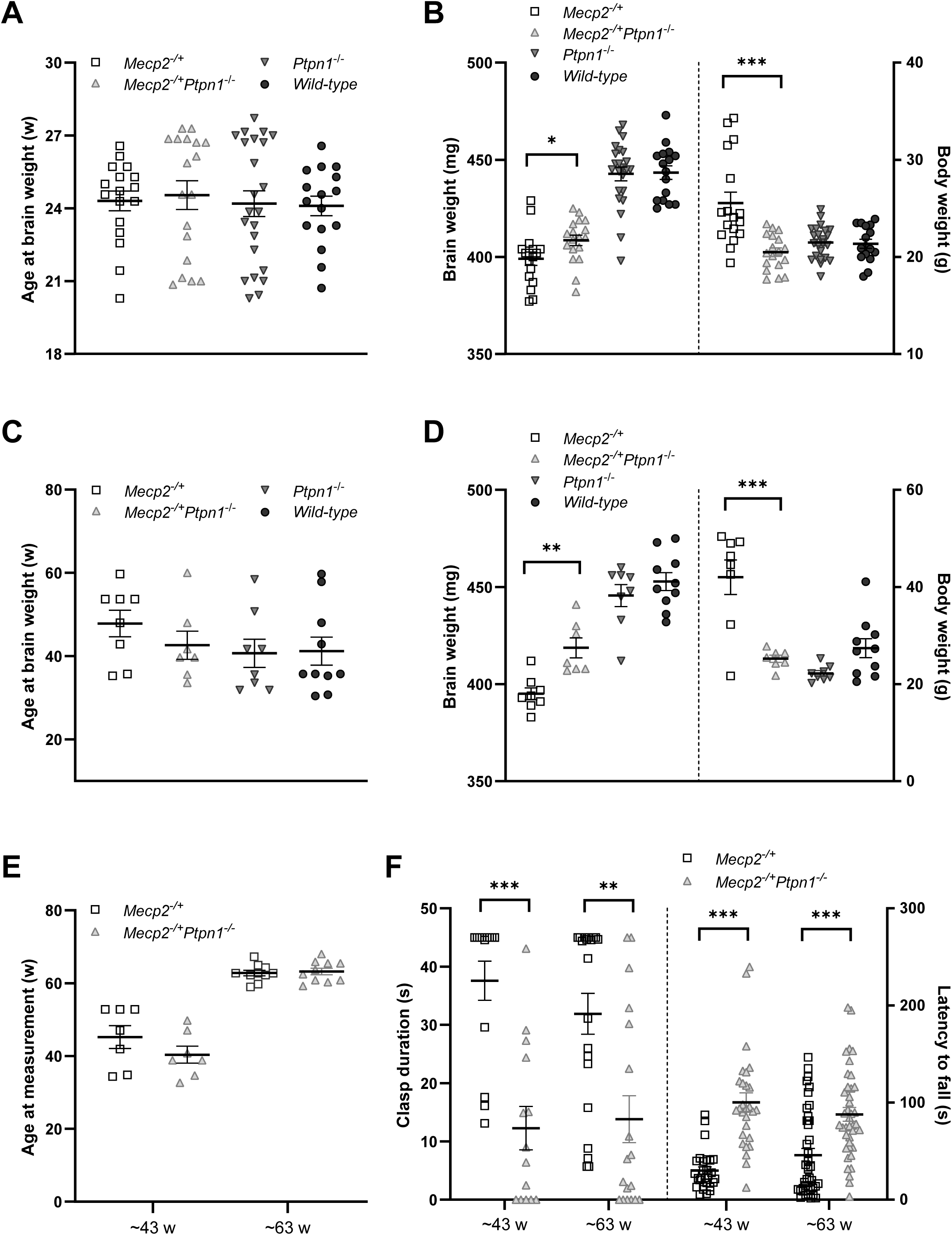
Loss of PTP1B increases brain weight and limits progression of motor and coordination deficits in *Mecp2^−/+^*mice. (**A** and **B**) Age and brain/body weight distribution measurements at ∼24 weeks of age (*Mecp2^−/+^*, *n* = 17; *Mecp2^−/+^Ptpn1^−/-^*, *n* = 18; *Ptpn1^−/-^*, *n* = 23; wild-type, *n* = 16). (**C** and **D**) Age and brain/body weight distribution measurements at ∼45 weeks of age (*Mecp2^−/+^*, *n* = 8; *Mecp2^−/+^Ptpn1^−/-^*, *n* = 7; *Ptpn1^−/-^*, *n* = 8; wild-type, *n* = 10). (**E** and **F**) Hind-limb clasp (range 0-45 seconds) and Rotarod performance (range 0-300 seconds) at later stages of disease progression, measured at ∼43 and ∼63 weeks of age, respectively. Clasping is expressed as the mean of two trials per animal, and latency to fall is expressed as the mean of four trials per animal (*Mecp2^−/+^*, *n* = 7; *Mecp2^−/+^Ptpn1^−/-^*, *n* = 7; *Mecp2^−/+^*, *n* = 10; *Mecp2^− /+^Ptpn1^−/-^*, *n* = 10). Data are presented as the mean ± SEM. Statistical significance was determined using unpaired *t*-tests * *p*< 0.05, ** *p*< 0.01, *** *p*< 0.001.

Longitudinal behavioral analysis demonstrated sustained improvements in motor function and hind-limb clasping over >50 weeks. In *Mecp2^−/+^* mice, the duration of clasping more than doubled (15 to 35 s) from 13 to 63 weeks of age (Fig. 3E, and Fig. 4, E & F), where nearly half of all trials (15 of 34) reached the maximum recording duration (45 s) at 43 to 63 weeks of age, thereby restricting the upper limit of the measurement (Fig. 4F). In contrast, the duration of clasping in *Mecp2^−/+^Ptpn1^−/-^*mice also doubled over the same time frame; however, clasping remained significantly less than in *Mecp2^−/+^* mice, increasing from ∼5 to 12 s (Fig. 3E and Fig. 4F). Overall, *Mecp2^⁻/+^Ptpn1^⁻/⁻^* mice maintained stable rotarod performance, whereas Mecp2^⁻/+^ mice exhibited progressive decline (Fig. 4F). Similarly, clasping severity remained significantly reduced across the lifespan. These data demonstrate that PTP1B inhibition produces durable, long-term correction of neurodevelopmental and functional deficits in RTT.

### Next-generation allosteric inhibitors demonstrate translational potential

To advance translational potential, we evaluated next-generation allosteric PTP1B inhibitors with improved pharmacological properties (Fig. 5A). Treatment of Mecp2^⁻/+^ mice with DPM-1003, an analogue of MSI-1436 [32,33], attenuated pathological weight gain and significantly improved motor performance over a 12-week period (Fig. 5, B–E). Unlike MSI-1436, DPM-1003 did not suppress normal weight gain in wild-type mice, suggesting an improved therapeutic window. Notably, treatment effects were partially reversible after drug withdrawal, with performance declining gradually but remaining improved relative to vehicle-treated controls (Fig. 5F and fig. S3A). This suggests that therapeutic benefit can be titrated and sustained with appropriate dosing strategies. These findings demonstrate that structurally distinct, allosteric inhibitors of PTP1B provide consistent therapeutic benefit and support further optimization for clinical application.

**Figure 5.**
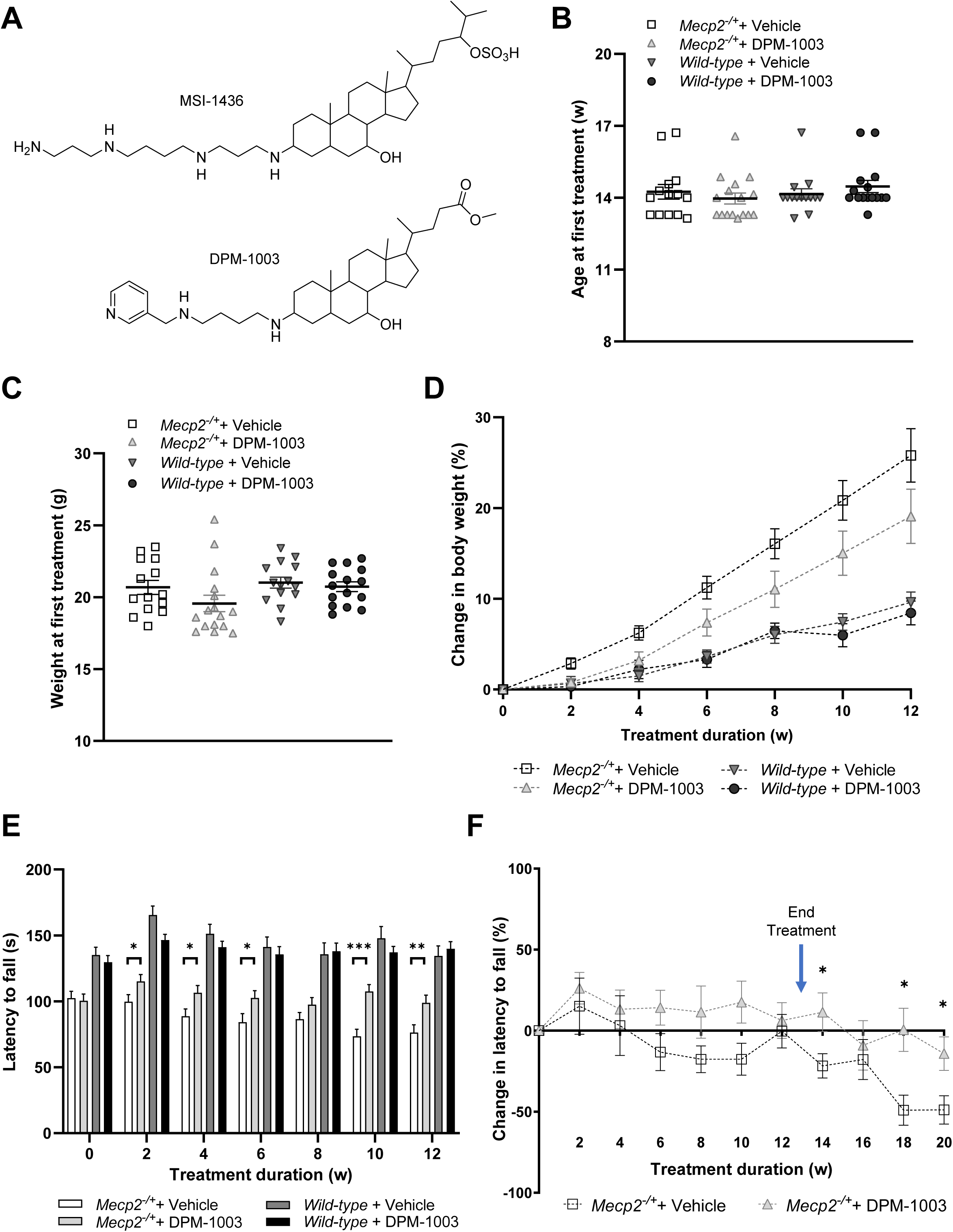
A next-generation allosteric PTP1B inhibitor reproduces therapeutic benefit in a mouse model of Rett Syndrome. (**A**) Chemical structures of allosteric PTP1B inhibitors MSI-1436 and DPM-1003. (**B**) Age distribution at initiation of treatment across cohorts (*Mecp2^−/+^* + vehicle, *n* = 14; *Mecp2^−/+^* + DPM-1003, *n* = 16; wild-type + Vehicle, *n* = 14; wild-type + DPM-1003, *n* = 15). (**C**) Body weight distribution at first treatment and (**D**) longitudinal change in body weight, expressed as the percent change in mean body weight per animal per week (*Mecp2^−/+^* + vehicle, *n* = 14; *Mecp2^−/+^* + DPM-1003, *n* = 16; wild-type + vehicle, *n* = 14; wild-type + DPM-1003, *n* = 15). (**E**) Rotarod performance, measured as latency to fall in a 0-300 second trial expressed as the mean of four trials per animal per week (*Mecp2^−/+^* + vehicle, *n* = 14; *Mecp2^−/+^* + DPM-1003, *n* = 16; wild-type + vehicle, *n* = 14; wild-type + DPM-1003, *n* = 15). (**F**) Change in Rotarod performance, expressed as percentage change in mean latency to fall from four trials per animal per week (*Mecp2^−/+^* + vehicle, *n* = 7; *Mecp2^−/+^*+ DPM-1003, *n* = 8). Data are presented as the mean ± SEM. Statistical significance was determined using unpaired *t*-tests * *p*< 0.05, ** *p*< 0.01, *** *p*< 0.001.

### PTP1B inhibition ameliorates cardio-respiratory dysfunction in an RTT mouse model

Cardio-respiratory dysfunction is a major contributor to morbidity and mortality in RTT. Sudden death of unknown cause occurs in ∼25% of all RTT patients, primarily during wakefulness [34,35]. Altered cardio-respiratory functions resulting from autonomic nervous system dysregulation is a major indicator of sudden death in RTT [36–38]. An important area of concern in RTT is exacerbation of cardiac arrythmias as result of prolonged corrected QT interval (QTc) caused by drug-induced cardiotoxicity or in combination with apnea-induced tachycardia. As Rett Syndrome patients have a predisposition to lethal cardiac arrythmias due to a prolonged QTc interval [39,40], we evaluated cardiac activity by non-invasive monitoring of electrocardiogram (ECG) signals in conscious *Mecp2^−/+^* mice prior to and post treatment with DPM-1003. Electrocardiographic analysis revealed prolonged QTc intervals in a subset of Mecp2^⁻/+^ mice (Fig. 6A), consistent with clinical observations in human subjects [39,40]. Treatment with DPM-1003 normalized QTc intervals in affected animals and reduced QTc across the cohort without evidence of cardiotoxicity (Fig. 6B), consistent with a cardioprotective effect.

**Fig. 6.**
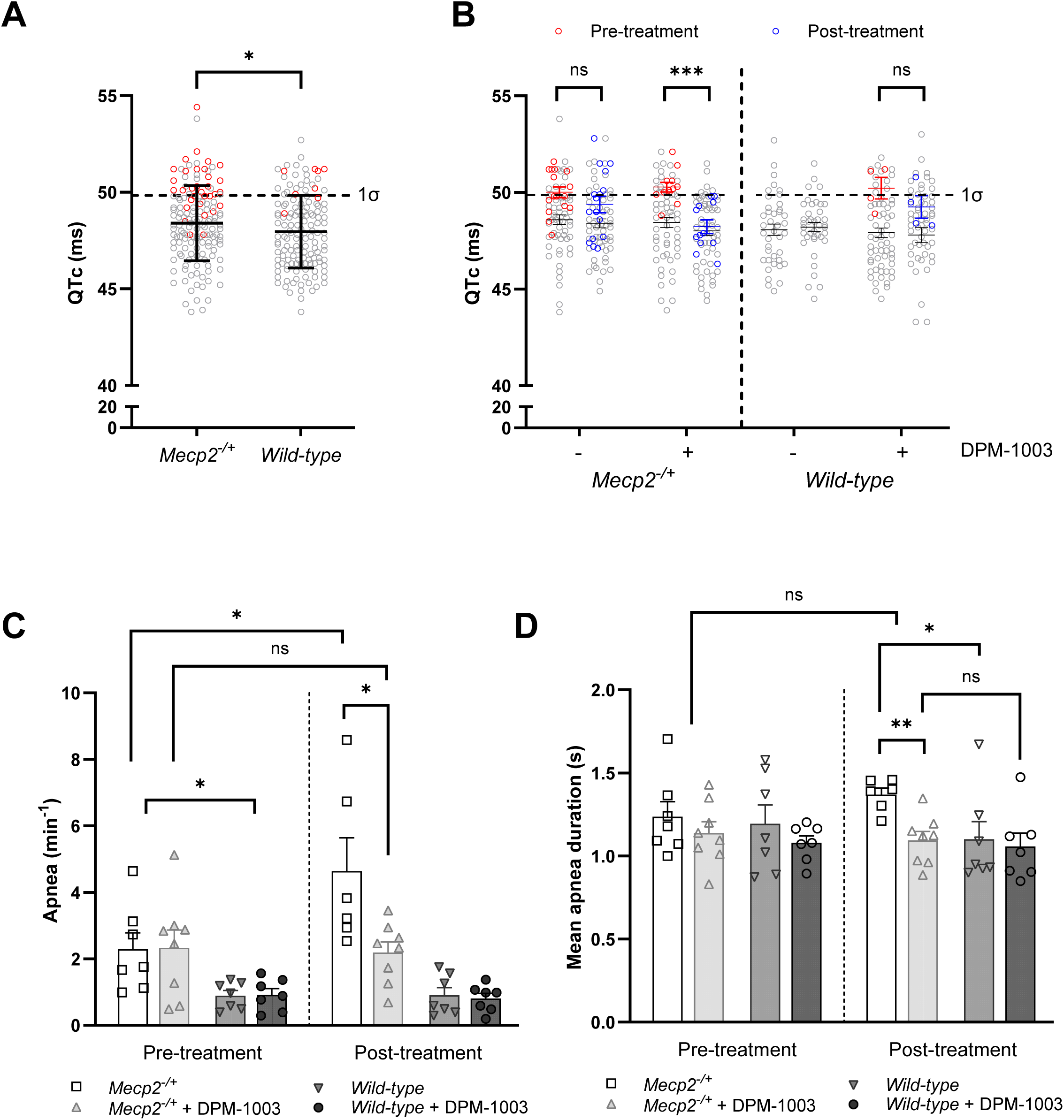
PTP1B inhibitor DPM-1003 ameliorates cardiac and respiratory dysfunction in a mouse model of Rett Syndrome. (**A**) Distribution of corrected QT (QTc) interval prior to the first treatment, calculated as the mean QTc interval from four electrocardiogram (ECG) traces per animal (*Mecp2^−/+^*, *n* = 37; wild-type, *n* = 35; from all traces). Prolonged QTc interval was defined as a pre-treatment four-trace (red) mean greater than one SD above the mean QTc interval of all traces from wild-type mice. Data represent the mean ± SD. (**B**) QTc intervals before and after treatment, calculated as the mean QTc interval from four ECG traces per animal (*Mecp2^−/+^*+ vehicle, *n* = 14; *Mecp2^−/+^* + DPM-1003, *n* = 14; wild-type + vehicle, *n* = 10; wild-type + DPM-1003, *n* = 15; colored in grey). Significance denotes comparison between animals exhibiting a prolonged QTc interval prior to treatment (*Mecp2^−/+^* + vehicle, *n* = 4; *Mecp2^− /+^* + DPM-1003, *n* = 3; wild-type + DPM-1003, *n* = 2; colored in red) with the same animal post-treatment (blue). Mice lacking four high-quality pre- or post-treatment ECG traces were excluded from analysis for that time point (*Mecp2^−/+^* + DPM-1003, *n* = 1, post-treatment; wild-type + vehicle, *n* = 2, post-treatment). None of the excluded mice met the predefined criteria for prolonged QTc. (**C**) Apnea frequency before and after treatment, expressed as the mean number of apneas per minute per animal (*Mecp2^−/+^* + vehicle, *n* = 7; *Mecp2^−/+^*+ DPM-1003, *n* = 8; wild-type + vehicle, *n* = 7; wild-type + DPM-1003, *n* = 7). (**D**) Mean apnea duration before and after treatment, calculated per animal in the same cohorts as in (C). Mice lacking observable apneas before or after treatment were excluded from analysis for that time point (*Mecp2^−/+^* + vehicle, *n* = 1, post-treatment). Data represent the mean ± SEM, unless otherwise indicated. Statistical significance was determined using unpaired *t*-tests * *p*< 0.05, ** *p*< 0.01, *** *p*< 0.001.

It has been reported that more than 80% of RTT patients experience breathing abnormalities throughout their lifespan that often manifest in the form of breath-holds [41–43]. Here, normal inspiration is followed by a brief post-inspiratory phase of decreased heart rate which is abruptly terminated by the onset of the breath-hold, concomitant with rapid tachycardia that can lead to severe cardio-respiratory complications. By contrast, breath-holds in healthy individuals are characterized by bradycardia (slowed heart rate) [44,45]. Animal models of RTT exhibit prolonged respiratory cycles and pauses during wakefulness (apneas) that are reminiscent of the clinical breath-hold phenotype [46]. Using whole-body plethysmography, we demonstrated increased apnea frequency and duration in *Mecp2^⁻/+^* mice. Treatment with DPM-1003 prevented worsening of apnea frequency and reduced apnea duration, restoring respiratory parameters to levels comparable to wild-type animals (Fig. 6, C & D). Together, these findings demonstrate that PTP1B inhibition improves core autonomic dysfunction in RTT, including both cardiac and respiratory phenotypes.

## DISCUSSION

Rett syndrome (RTT), affecting ∼1 in 10,000 individuals, is a profoundly challenging neurodevelopmental disorder in which affected children initially undergo a period of apparently normal development, followed by rapid and sustained neurological regression beginning at 6–18 months of age. Early manifestations often include autistic-like features, such as social withdrawal and repetitive behaviors, but the disease evolves into a multisystem disorder characterized by severe impairments in motor coordination, seizures, and cardiorespiratory and gastrointestinal dysfunction [2–5]. The breadth and progression of these phenotypes underscore the need for therapeutic strategies that address core disease biology rather than isolated symptoms.

The recent approval of Trofinetide represents an important milestone for the field [47–50]. In clinical studies employing the Rett Syndrome Behavioral Questionnaire (RSBQ) and Clinical Global Impression–Improvement (CGI-I) scales, Trofinetide demonstrated modest improvements in communication, engagement, and motor function; however, its clinical utility is constrained by a high incidence of gastrointestinal adverse events. In the Phase 3 LAVENDER study, diarrhea was reported in >80% of patients, with over half requiring antidiarrheal intervention; furthermore, the Phase 2/3 DAFFODIL study similarly reported substantial rates of vomiting. These tolerability issues, likely linked to the relatively high doses required to achieve clinical benefit, present practical challenges for long-term treatment and patient adherence [51–54].

Trofinetide is derived from the N-terminal tripeptide (Gly–Pro–Glu) of IGF-1 [55–58], prompting initial hypotheses that it’s mechanism of action may be through modulation of IGF-1 receptor signaling. Nevertheless, structural and biochemical studies do not support direct engagement of the IGF-1 receptor by this region of the ligand, and preclinical binding assays have failed to identify a definitive molecular target [59]. Consistent with this uncertainty, Trofinetide has shown limited efficacy across several preclinical endpoints relevant to RTT pathophysiology, including motor performance and respiratory function. Thus, although Trofinetide provides symptomatic benefit in some patients, its mechanism of action remains incompletely defined, and it does not appear to address directly the underlying molecular defects associated with MECP2 dysfunction.

Taken together, these observations highlight a critical gap in the current therapeutic landscape; the absence of mechanism-based interventions that target the core signaling pathways disrupted in RTT downstream of MECP2 dysfunction. This gap provides a strong rationale for the development of next-generation therapies grounded in disease biology, with the potential to achieve more durable and system-wide clinical benefit. In this study, we demonstrate that inhibition of the protein tyrosine phosphatase PTP1B represents a mechanism-based therapeutic strategy capable of alleviating a broad range of pathological manifestations in female murine models of RTT. Both pharmacological inhibition, using brain-penetrant allosteric small molecules, and genetic ablation of PTP1B produced consistent improvements in motor performance, neuromuscular function, neurodevelopmental deficits, and cardio-respiratory abnormalities. These findings provide compelling evidence that PTP1B is a disease-relevant signaling node in RTT and validate its inhibition as a promising therapeutic approach for modifying disease pathology rather than alleviating symptoms alone.

Our findings build on previous work demonstrating that MECP2 directly represses transcription of the PTPN1 gene, which encodes PTP1B [16]. Loss of MECP2 function leads to elevated PTP1B expression, thereby imposing a barrier to neurotrophic signaling pathways that are essential for normal neuronal function. In particular, signaling through the BDNF receptor TRKB has been implicated as a critical pathway that is disrupted in RTT. Rather than attempting to stimulate TRKB directly through receptor agonists, which may bypass physiological regulatory mechanisms, inhibition of PTP1B provides an alternative strategy to overcome the barrier to signaling and restore endogenous responsiveness to neurotrophic stimulation. This approach may offer advantages in preserving physiological signaling dynamics while overcoming the inhibitory constraints imposed by increased phosphatase activity.

An important feature of the present study is the concordance between pharmacological and genetic approaches to Inhibition of PTP1B function. Deletion of PTP1B in *Mecp2*-mutant mice reproduced many of the phenotypic improvements observed with pharmacological inhibition, including restoration of motor coordination, reduction in hind-limb clasping, increased grip strength, and attenuation of excessive weight gain. The agreement between these independent strategies strongly supports the conclusion that the therapeutic effects observed arise from on-target inhibition of PTP1B rather than off-target drug activity. Moreover, the absence of phenotypic alterations in wild-type animals lacking PTP1B further suggests that the improvements observed in RTT models reflect restoration of MECP2-dependent physiological processes rather than generalized enhancement of motor or metabolic function.

The breadth of phenotypic rescue observed following PTP1B inhibition is particularly noteworthy. Rett syndrome affects multiple neural circuits governing motor control, autonomic regulation, and neurodevelopment, and therapeutic approaches that address only a single symptom or pathway have had limited success. In this context, the improvements observed in motor coordination, muscle strength, microcephaly, cardiac electrophysiology, and respiratory function suggest that PTP1B inhibition acts at a systems level to restore signaling pathways that are broadly dysregulated by MECP2 deficiency. The localization of PTP1B inhibitors to regions of the brain involved in motor coordination and autonomic regulation, including the medial prefrontal cortex and brainstem, further supports the notion that pharmacological inhibition of PTP1B engages neural circuits that are central to the pathophysiology of Rett syndrome.

Another notable aspect of our findings is the durability of the therapeutic effects observed. Improvements in behavioral and physiological phenotypes were sustained over extended periods in both pharmacological and genetic models, with benefits persisting for many months. Similar observations have been made in response to pre-symptomatic training, which together with PTP1B inhibition, may harbor additional restorative benefits and broader therapeutic windows [60]. Such long-term improvements suggest that modulation of signaling pathways downstream of MECP2 may influence developmental or circuit-level processes that extend beyond transient pharmacological effects. This possibility is supported by the observation that PTP1B ablation partially restores brain weight in *Mecp2*-heterozygous mice, consistent with previous studies showing that reduced brain size in RTT reflects impaired neuronal maturation rather than neurodegeneration [27–31]. Restoration of neurotrophic signaling pathways may therefore contribute to improved neuronal connectivity and circuit stability over time.

From a translational perspective, our study highlights the therapeutic potential of selectively targeting a protein tyrosine phosphatase in neurological disease [21]. Historically, phosphatases have been considered challenging drug targets due to the conserved and highly charged nature of their catalytic sites. The development of small molecule allosteric inhibitors of PTP1B, which bind at sites distinct from the catalytic pocket, has provided an opportunity to overcome these challenges. The compounds used in this study are specific for PTP1B, orally bioavailable, cross the blood–brain barrier, and demonstrate clear pharmacodynamic activity in vivo. Importantly, pharmacological inhibition of PTP1B ameliorated QT interval abnormalities and respiratory disturbances in RTT mouse models, suggesting that targeting this pathway may address life-threatening autonomic complications that contribute to morbidity and mortality in patients.

Current therapeutic strategies for RTT include gene replacement or gene editing approaches aimed at restoring MECP2 expression. Although these approaches hold promise, they face significant challenges related to dosage sensitivity, mosaic expression, and delivery to the central nervous system. Both insufficient and excessive MECP2 expression can produce severe neurological phenotypes, therefore achieving precise regulation of MECP2 levels remains a critical barrier to clinical implementation [11,12]. In contrast, modulation of downstream signaling pathways through small molecule drugs may provide a complementary strategy that bypasses these challenges while still addressing key molecular consequences of MECP2 deficiency.

Although the present study establishes PTP1B inhibition as a mechanism-based strategy that restores signaling and improves multiple disease-relevant phenotypes in RTT models, future studies will be important to define its long-term efficacy and translational potential in patients. Furthermore, although the phenotypic improvements observed are substantial, the precise molecular mechanisms linking PTP1B inhibition to restoration of neuronal and autonomic function remain to be fully elucidated. Enhanced TRKB signaling is likely to play a central role, however, PTP1B regulates multiple signaling pathways, including those involved in metabolic regulation and neuronal plasticity. Future studies aimed at defining the signaling networks and cellular targets affected by PTP1B inhibition in specific neuronal populations will provide important insights into the mechanisms underlying therapeutic benefit.

In summary, our findings establish PTP1B as a mechanistically grounded and therapeutically actionable target in Rett syndrome. Pharmacological inhibition of PTP1B restores signaling pathways disrupted by MECP2 deficiency and produces broad and sustained improvements in neurological and autonomic phenotypes in physiologically relevant models of the disease. These results support a new therapeutic paradigm in which selective modulation of signal transduction pathways can compensate for genetic defects in transcriptional regulation and provide meaningful clinical benefit. The recent clearance of an Investigational New Drug application for clinical testing of PTP1B inhibitors further underscores the translational potential of this strategy and provides a strong rationale for evaluating PTP1B inhibition as a disease-modifying therapy in patients with Rett syndrome.

## ACKNOWLEDGEMENTS

NKT is the Caryl Boies Professor of Cancer Research at Cold Spring Harbor Laboratory. We thank R. Rubino, L. Bianco, J. Habel, and staff at the Animal Shared Resource of the Cold Spring Harbor Laboratory Cancer Center for animal procedure, housing, and maintenance support. We are grateful to A. Grill (DepYmed Inc.) for providing PTP1B inhibitors, B. D. Ross (University of Michigan) for use of computational resources and research infrastructure, T. Hampton (Mouse Specifics, Inc.) for assistance with electrocardiogram data analysis, and L. Bogdanik and Z. Beaudry (The Jackson Laboratory) for assistance with plethysmography data analysis.

## Author contributions

Conceptualization: NKT, CAB

Methodology and data analysis: CAB, CF, L.N.C

Funding acquisition and Resources: NKT

Project administration: NKT

Supervision: NKT and CAB

Writing: NKT and CAB with review and editing by all authors.

All authors have approved the submitted version of the manuscript

## Data and materials availability

Data generated during this study are available from the corresponding author on reasonable request. All data for evaluation of conclusions are present in the paper or the Supplementary Materials.

## MATERIALS AND METHODS

### Animals

Mice used in this study were housed in the animal vivarium at Cold Spring Harbor Laboratory. All experimental protocols were reviewed and approved by the Cold Spring Harbor Laboratory Institutional Animal Care and Use Committee (IACUC; Protocol number 2024-1382) and were conducted in accordance with the NIH’s Guide for the Care and Use of Laboratory Animals. Wild-type C57BL/6J mice (wild-type), B6.129P2(C)-Mecp2^tm1.1Bird^/J mice (*Mecp2^−/+^*), and B6.129S4-*Ptpn1^tm1Bbk^*/Mmjax mice (*Ptpn1^−/-^*) were obtained from The Jackson Laboratory. Male B6.129S4-*Ptpn1^tm1Bbk^*/Mmjax (*Ptpn1^−/-^*) and C57BL/6J (wild-type) mice were bred in-house to female C57BL/6J (wild-type) and B6.129P2(C)-Mecp2^tm1.1Bird^/J (*Mecp2^−/+^*) mice to generate the following offspring for behavioral experiments, strain and breeding colony generation, and maintenance: *Mecp2^−/+^*, *Mecp2^−/+^Ptpn1^−/+^*, *Mecp2^−/+^Ptpn1^−/-^*, *Ptpn1^−/+^*, *Ptpn1^−/-^*, and wild-type female mice, and *Ptpn1^−/-^* and wild-type male mice. Mice were housed up to five per cage and maintained on a 12h light/dark cycle at ambient temperature (25 °C), with standard murine chow and water *ad libitum*.

### Compound administration

Compounds (MSI-1436 and DPM-1003; DepYmed Inc.) were dissolved in sterile normal saline solution (Vehicle) and administered twice weekly as a single intraperitoneal injection at 2 mg/kg for 6 weeks (MSI-1436) or 5 mg/kg for 12 weeks (DPM-1003), beginning at a mean age of 12-14 weeks. For pharmacokinetic studies (WuXi AppTec Corp.), compound (MSI-1436) was dissolved in sterile phosphate buffered saline (PBS) and administered as a single intraperitoneal or intravenous injection at 2 mg/kg to male C57BL/6J (*Wild-type*) mice. At designated time points, blood plasma (EDTA-K2) or brain tissue homogenates were collected, then protein was precipitated and compound isolated by methanol extraction containing internal standards (Labetalol, Tolbutamide, Diclofenac; 100 ng/ml) at a ratio of 1:6 to 1:8. Sample extracts were further mixed ∼2:1 with water, and analyzed by reverse-phase LC-MS/MS against a matrix-matched 8-point standard curve. Concentration measurements > 50-fold less than the mean of all other animals, over multiple time points, were suspected to have been compromised by handler error and were excluded from analysis. For mass spectrometry imaging studies, compound (MSI-1436) was dissolved in sterile normal saline solution (Vehicle) and administered twice weekly as a single intraperitoneal injection at 10 mg/kg for 2 w to *Mecp2^−/+^* and *Wild-type* mice at an average age of 22 weeks. Animals were euthanatized 2-3 h after final compound administration (Tmax), brains were removed by dissection, frozen on dry ice, then transferred to -80 °C. Fresh-frozen brain tissue was sectioned (sagittal) in a cryostat (Leica CM1950) maintained at -18 °C until ∼2 mm from the mid-line, then serial 10 µm sections were thaw mounted onto glass microscope slides (FisherScientific), and slides desiccated under vacuum for ≥ 20 min at ambient temperature, then transferred to a slide box in a sealable bag containing desiccant packs and stored at -80 °C until analysis. Slides were brought to ambient temperature under vacuum desiccation, then coated with 2,5-dihydroxybenzoic acid (Sigma) in 50% methanol 0.1% trifluoroacetic acid (FisherScientific) using an HTX M5 Sprayer (HTX Technologies), and analyzed by mass spectrometry imaging on a Waters Synapt G2Si MALDI-TOF mass spectrometer using Waters High-Definition Imaging (HDI) and MassLynx software platforms (Waters Corp.) for data acquisition and processing. Purified compound (MSI-1436) was also spotted onto a MALDI target plate and adjacent tissue as a reference mass and standard for comparison to compound distribution in treated animals.

### Study design

#### Acclimatization

Prior to collecting experimental measurements, animals undergo an acclimatization period (∼2 weeks) that includes saline injections (intraperitoneally; twice weekly) and/or body weight measurements, and behavioral training to gain familiarity with routine handling, restraint, compound administration, and the experimental apparatus. After this time, pre-treatment or baseline physical, behavioral, and physiological assessments are measured, then studies initiated. Animals are routinely acclimatized for ∼30 min following transport out of general housing area to the behavioral space, and instruments cleaned with disinfectant wipes prior to recording of any measurements, and before, after, and in-between subjects.

#### Animal cohorts and grouping

Animal cohorts were separated based on genotype, gender, body weight, rotarod performance, and litter to establish pre-treatment or baseline groupings with equivalent mean body weight and variance, latency to fall, and representative distribution of littermates, among all cohort groups over the study duration. Rotarod, which measures motor coordination and balance, was chosen as a primary criterion for its quantitation and reproducibility.

#### Weight measurement

Animal body weight is measured at least twice weekly prior to compound administration and/or behavioral assessment and is calculated as the weekly average of two measurements. Animals are monitored regularly for overall health, locomotion, social interactions, eating, drinking, and grooming behavior, and for excessive weight loss/fluctuations and malocclusion. When identified, animals are provided access to Diet Gel® and/or moistened food in the cage bottom, teeth are clipped, and data is curated to remove animals where body weight accuracy may have been compromised as result of non-experimental factors. Animals missing equivalent pre-treatment, baseline, or weekly body weight measurements are excluded from analysis or for that time point, respectively. Animal brain weight is measured at time of sacrifice, after inspection for gross abnormalities or atrophy.

#### Paw clasping

Hind-limb clasping is measured once weekly within ∼24-48 h of compound administration or target age range and is calculated from the duration of hind-limb clasping observed over a 45 s period from two separate trials. We have defined a clasp as the retraction of one or both of the hind limbs when the animal is suspended by their tail, such that the paw of the retracted limb is drawn in to the animals’ body and the toes are curled into a fist shape. Hind limbs or paws do not need to touch to be considered a clasp by this definition. Animals missing equivalent measurements were excluded from analysis for that time point.

#### Rotarod

Latency to fall is measured once weekly within ∼24 hours of compound administration or target age range and is calculated from the duration of time spent on an increasing angular speed rotating rod (4-40 rpm) over a maximum 300 seconds (5 min) period in four separate trials. Animals undergo initial acclimatization training at a static angular speed (4 rpm) for 300 seconds, where fallen mice are placed back on the rotating rod and corrected if begin walk backwards. This is followed by a 10 min rest, then four trials of increasing angular speed (4-40 rpm) for 300 seconds, each with a 10 min rest between trials. The same regimen is performed 24 hours later to obtain pre-treatment or baseline measurements, then continued weekly or at defined ages until study end point. Data is curated to exclude individual trials or animals where latency to fall may have been the result of environmental factors (causing trial latency < 50% of next lowest trial), instrument malfunction, or identification as outliers by the Rout method (Q = 1%). Animals missing equivalent measurements or with measurements repeatedly identified as outliers over multiple weeks were excluded for that time point or from the analysis, respectively. By comparison, data curation had no effect on the overall observations or conclusions of the study.

#### Grip strength

Front-limb grip strength is measured within ∼24 hours of compound administration or target age range and is calculated from the maximum resistance (grams of force) of symmetrical front-limb grip (both paws) to the apparatus upon application of an opposing force until release, from five separate trials. Prior to measurement, the metering apparatus is monitored for consistency by taking several readings of the force required to break contact between a paperclip/magnet pair. Animals undergo initial acclimatization training 48 hour prior to experimental data acquisition, and longitudinal testing is repeated no more than once every two weeks to avoid habituation. Mice are gently held by the tail and lowered until front paws are at the same height as a trapeze bar attached to the metering device that is parallel to the benchtop, and a firm grip is established. Mice are then positioned horizontally to the apparatus, and a smooth, steady pulling motion is applied until grip is broken. Force values are recorded, and this is repeated up to 5 times with a 10 min rest between each trial.

#### Electrocardiogram

Electrical activity of the heart is measured continuously for 5-10 min in conscious, unrestrained mice within 2-3 h of compound administration (Tmax), and functional cardiac metrics and heart rate variability indices are calculated from an ensemble average electrocardiogram waveform (ECGenie; Mouse Specifics, Inc.). Animals are placed on an elevated mesh platform connected to the system for ∼10 min to acclimatize, then allowed to freely move into the recording chamber platform by way of a sliding door to avoid handling stress. Measurements are recorded and monitored in real-time for signal quality to obtain ≥ 4 high-quality, non-invasive ECG traces of sufficient length from animals that are still, with paws making adequate electrode contact. Raw data is analyzed using the EzCG software, with baseline drift and signal artifacts associated with animal movement removed manually (guided by prompts in the software) to automatically generate the ensemble average electrocardiogram waveform and tabulate all ECG data processing metrics and indices including corrected QT interval (QTc) for each trace (4 per animal). We have defined a prolonged QTc interval phenotype as any animal displaying a pre-treatment four-trace average QTc interval greater than one standard deviation (1 σ) of the mean QTc interval of all traces from wild-type mice. Animals without ≥ 4 high-quality ECG traces are excluded from analysis. No excluded animals exhibited the prolonged QTc interval phenotype as defined.

#### Plethysmography

Respiratory activity is measured continuously for 3 hours in conscious, unrestrained mice within ∼24 hours of compound administration by whole-body plethysmography (Buxco WBP; Data Sciences International, DSI), with functional respiratory metrics and breathing cycle indices calculated from a collection of stand-still periods (SSPs) acquired using the FinePointe Respiratory and Inhalation software (DSI). Animals (8) are placed into individual acrylic WBP chambers connected to the system with electronically regulated airflow for ∼30 min to acclimatize prior to initiating data acquisition, being monitored during this time for signs of distress. Measurements are recorded and analyzed for signal quality to obtain ≥ 10 min of high-quality, non-invasive SSPs of adequate length (≥ 3 min each) from awake animals that are still and non-exploritory (not grooming or sniffing). Raw data is analyzed using the Plethyproc software (The Jackson Laboratory) to identify individual breathing cycles from the start of inspiration of a given cycle to the start of inspiration of the following cycle for calculataion of the duration of inspiration (Ti), duration of expiration (Te), and duration of the total breathing cycle (Ttot). We have defined an apnea as Te > 2 * Ttot for calculation of the number of apneas per minute and the mean apnea duration. Animals without any observed pre- or post-treatment apneas were excluded from analysis for that time point.

### Statistical Analysis

Statistical analysis was performed using GraphPad Prism 10. All data representations and statistical tests are specified in the figure legends. Data are presented as mean ± SEM or SD, with significance evaluated using two-tailed, unpaired Student’s *t*-test with a threshold of α = 0.05 (ns *p* > 0.05, * *p* < 0.05, ** *p* < 0.01, *** *p* < 0.001). Outliers were evaluated by the Rout method (Q = 1%). Animal cohort and sample sizes were determined on the basis of prior experience and data characterizing the studied phenotypes in B6.129P2(C)-Mecp2^tm1.1Bird^/J and B6.129S4-*Ptpn1^tm1Bbk^*/Mmjax mice (*Mecp2^−/+^* and *Ptpn1^−/-^*, respectively).

## SUPPLEMENTARY FIGURE LEGENDS

**Figure S1. Retrospective analysis demonstrates consistency and reproducibility of the experimental platform across cohorts.** (**A**) Age distribution at first treatment or first measurement across all cohorts (*Mecp2^−/+^* + vehicle, *n* = 34; *Mecp2^−/+^* + PTP1B inhibitor, *n* = 38; wild-type + Vehicle, *n* = 32; wild-type + PTP1B inhibitor, *n* = 37; *Mecp2^−/+^*, *n* = 43; wild-type, *n*=34).

(**B**) Body weight distribution at first treatment or first measurement across all cohorts (*Mecp2^−/+^* + Vehicle, *n* = 33; *Mecp2^−/+^* + PTP1B inhibitor, *n* = 38; wild-type + vehicle, *n* = 31; wild-type + PTP1B inhibitor, *n* = 36; *Mecp2^−/+^*, *n* = 39; wild-type, *n*=30). Mice lacking equivalent body weight measurements were excluded from analysis for that time point. Mice with repeated body weight measurement fluctuations attributed to malocclusion were excluded from analysis.

(**C**) Rotarod performance at first treatment or first measurement, expressed as latency to fall from four trials per animal across all cohorts (*Mecp2^−/+^* + vehicle, *n* = 34; *Mecp2^−/+^* + PTP1B inhibitor, *n* = 38; wild-type + vehicle, *n* = 32; wild-type + PTP1B inhibitor, *n* = 37; *Mecp2^−/+^*, *n* = 42; wild-type, *n*=34; range, 0-300 seconds). Mice with measurements repeatedly identified as outliers over multiple weeks were excluded from analysis.

(**D**) Rotarod performance at first treatment or first measurement, expressed as the four-trial mean across all cohorts (same cohorts as in C).

A-D represents a compilation of data shown in Figures 1B and C, 2B, 3B and D, and 5B, C, and E.

(**E**) Plasma pharmacokinetics of MSI-1436 following a single intravenous injection per animal, shown as the mean blood plasma concentration at each time point (wild-type + MSI-1436, 0.5 mg/kg, *n* = 6; wild-type + MSI-1436, 2 mg/kg, *n* = 6).

(**F**) Hind-limb clasping at first treatment, expressed as mean clasp duration from two trials per animal across three cohorts (*Mecp2^−/+^* + vehicle, *n* = 19; *Mecp2^−/+^* + MSI-1436, *n* = 22; wild-type + vehicle, *n* = 6; wild-type + MSI-1436, *n* = 8; range 0-45 seconds). Mice missing equivalent measurements were excluded from analysis. These data are a compilation from experiments shown in Figure 2C.

Together, these analyses support the internal consistency and reproducibility of the phenotyping platform used across pharmacological and genetic studies. Data are presented as mean ± SEM. Outliers were evaluated by the Rout method (Q = 1%). Statistical significance was determined using unpaired *t*-tests * *p*< 0.05, ** *p*< 0.01, *** *p*< 0.001.

**Figure S2. Genetic reduction or loss of PTP1B improves physical and behavioral phenotypes in *Mecp2^−/+^* mice.** (**A**) Age distribution at first treatment across genotypes (*Mecp2^−/+^*, *n* = 18; *Mecp2^−/+^Ptpn1^−/+^*, *n* = 26; *Mecp2^−/+^Ptpn1^−/-^*, *n* = 24; *Ptpn1^−/+^*, *n* = 32; *Ptpn1^−/-^*, *n* = 23; wild-type, *n* = 16).

(**B**) Body weight distribution at first measurement and (**C**) longitudinal change in body weight, expressed as the percent change in body weight per animal per week (*Mecp2^−/+^*, *n* = 17; *Mecp2^− /+^Ptpn1^−/+^*, *n* = 24; *Mecp2^−/+^Ptpn1^−/-^*, *n* = 24; *Ptpn1^−/+^*, *n* = 31; *Ptpn1^−/-^*, *n* = 23; wild-type, *n* = 16). Significance denotes comparison between *Mecp2^−/+^* and *Mecp2^−/+^Ptpn1^−/-^* (top), and between wild-type and *Mecp2^−/+^Ptpn1^−/-^*(bottom). Mice lacking equivalent body weight measurements were excluded from analysis for the indicated time point (*Mecp2^−/+^*, *n* = 3, Week 3; *Mecp2^−/+^Ptpn1^−/+^*, *n* = 6, Week 3; *Mecp2^−/+^Ptpn1^−/-^*, *n* = 5, Week 3; *Ptpn1^−/+^*, *n* = 2, Week 3; *Ptpn1^−/-^*, *n* = 4, Week 3; *Wild-type*, *n* = 2, Week 3). Mice with repeated body weight fluctuations attributed to malocclusion were excluded from analysis (*Mecp2^−/+^*, *n* = 1; *Mecp2^−/+^Ptpn1^−/+^*, *n* = 2; *Ptpn1^−/+^*, *n* = 1).

(**D**) Rotarod performance, measured as latency to fall and expressed as the mean of four trials per animal per week (*Mecp2^−/+^*, *n* = 17; *Mecp2^−/+^Ptpn1^−/+^*, *n* = 25; *Mecp2^−/+^Ptpn1^−/-^*, *n* = 24;*Ptpn1^−/+^*, *n* = 32; *Ptpn1^−/-^*, *n* = 23; wild-type, *n* = 16; range 0-300 seconds). Mice with measurements repeatedly identified as outliers over multiple weeks were excluded from analysis (*Mecp2^−/+^*, *n* = 1; *Mecp2^−/+^Ptpn1^−/+^*, *n* = 1).

(B-D) These data extend Figures 3B-D by including mice heterozygous for *Ptpn1* (*Ptpn1^−/+^*) and further support a gene dosage-dependent contribution of PTP1B to disease-relevant phenotypes in Mecp2^+/−^ mice.

(**E** and **F**) Age and brain/body weight measurements at ∼63 weeks of age across cohorts (*Mecp2^− /+^*, *n* = 10; *Mecp2^−/+^Ptpn1^−/-^*, *n* = 9; *Ptpn1^−/-^*, *n* = 5; wild-type, *n* = 2).

Data are presented as mean ± SEM. Outliers were evaluated by the Rout method (Q = 1%). Statistical significance was determined using unpaired *t*-tests * *p*< 0.05, ** *p*< 0.01, *** *p*< 0.001.

**Fig. S3. Behavioral benefit of DPM-1003 is consistent with reversible allosteric inhibition of PTP1B.** Change in Rotarod performance, expressed as percent change in mean latency to fall from four trials per animal per week (*Mecp2^−/+^* + vehicle, *n* = 7; *Mecp2^−/+^*+ DPM-1003, *n* = 8; wild-type + vehicle, *n* = 7; wild-type + DPM-1003, *n* = 8; range 0-300 seconds). Significance denotes comparison between *Mecp2^−/+^* + vehicle and *Mecp2^−/+^* + DPM-1003. These data extend Fig. 5F by including wild-type animals treated with vehicle or DPM-1003 and support the conclusion that reversible allosteric inhibition of PTP1B is sufficient to improve motor performance in Mecp2+/− mice.

Data are presented the mean ± SEM. Statistical significance was determined using unpaired *t*-tests * *p*< 0.05, ** *p*< 0.01, *** *p*< 0.001.

## REFERENCES

1. Amir, R.E., Van den Veyver, I.B., Wan, M., Tran, C.Q., Francke, U., and Zoghbi, H.Y. (1999). Rett syndrome is caused by mutations in X-linked MECP2, encoding methyl-CpG-binding protein 2. Nat Genet 23, 185–188.

2. Lopes, A.G., Loganathan, S.K., and Caliaperumal, J. (2024). Rett Syndrome and the Role of MECP2: Signaling to Clinical Trials. Brain Sci 14. 10.3390/brainsci14020120.

3. Gold, W.A., Percy, A.K., Neul, J.L., Cobb, S.R., Pozzo-Miller, L., Issar, J.K., Ben-Zeev, B., Vignoli, A., and Kaufmann, W.E. (2024). Rett syndrome. Nat Rev Dis Primers 10, 84.

4. Percy, A.K. (2011). Rett syndrome: exploring the autism link. Arch Neurol 68, 985–989.

5. Bricker, K., and Vaughn, B.V. (2024). Rett syndrome: a review of clinical manifestations and therapeutic approaches. Frontiers in Sleep Volume 3-2024. 10.3389/frsle.2024.1373489.

6. Cheval, H., Guy, J., Merusi, C., De Sousa, D., Selfridge, J., and Bird, A. (2012). Postnatal inactivation reveals enhanced requirement for MeCP2 at distinct age windows. Hum Mol Genet 21, 3806–3814.

7. McGraw, C.M., Samaco, R.C., and Zoghbi, H.Y. (2011). Adult neural function requires MeCP2. Science 333, 186. 10.1126/science.1206593.

8. Nguyen, M.V., Du, F., Felice, C.A., Shan, X., Nigam, A., Mandel, G., Robinson, J.K., and Ballas, N. (2012). MeCP2 is critical for maintaining mature neuronal networks and global brain anatomy during late stages of postnatal brain development and in the mature adult brain. J Neurosci 32, 10021–10034.

9. Guy, J., Gan, J., Selfridge, J., Cobb, S., and Bird, A. (2007). Reversal of neurological defects in a mouse model of Rett syndrome. Science 315, 1143–1147.

10. Alvarez-Saavedra, M., Saez, M.A., Kang, D., Zoghbi, H.Y., and Young, J.I. (2007). Cell-specific expression of wild-type MeCP2 in mouse models of Rett syndrome yields insight about pathogenesis. Hum Mol Genet 16, 2315–2325.

11. Lombardi LM, Baker SA, Zoghbi HY. (2015) MECP2 disorders: from the clinic to mice and back. Journal of Clinical Investigation. 125: 2914–2923.

12. Ta, D., Downs, J., Baynam, G., Wilson, A., Richmond, P., and Leonard, H. (2022). A brief history of MECP2 duplication syndrome: 20-years of clinical understanding. Orphanet J Rare Dis 17, 131. 10.1186/s13023-022-02278-w.

13. Leonard, H., Cobb, S., and Downs, J. (2017). Clinical and biological progress over 50 years in Rett syndrome. Nat Rev Neurol 13, 37–51. 10.1038/nrneurol.2016.186.

14. Percy, A.K., Ananth, A., and Neul, J.L. (2024). Rett Syndrome: The Emerging Landscape of Treatment Strategies. CNS Drugs 38, 851–867. 10.1007/s40263-024-01106-y.

15. Adams, I., Yang, T., Longo, F.M., and Katz, D.M. (2020). Restoration of motor learning in a mouse model of Rett syndrome following long-term treatment with a novel small-molecule activator of TrkB. Dis Model Mech 13. 10.1242/dmm.044685.

16. Krishnan, N., Krishnan, K., Connors, C.R., Choy, M.S., Page, R., Peti, W., Van Aelst, L., Shea, S.D., and Tonks, N.K. (2015). PTP1B inhibition suggests a therapeutic strategy for Rett syndrome. J Clin Invest 125, 3163–3177.

17. Ricceri L, De Filippis B, Laviola G. (2008) Mouse models of Rett syndrome: from behavioural phenotyping to preclinical evaluation of new therapeutic approaches. Behavioural Pharmacology. 19: 501–517.

18. Allemang-Grand, R., Ellegood, J., Spencer Noakes, L., Ruston, J., Justice, M., Nieman, B.J., and Lerch, J.P. (2017). Neuroanatomy in mouse models of Rett syndrome is related to the severity of Mecp2 mutation and behavioral phenotypes. Mol Autism 8, 32.

19. Katz DM, Bird A, Coenraads M, Gray SJ, Menon DU, Philpot BD, Tarquinio DC. (2016) Rett Syndrome: Crossing the Threshold to Clinical Translation. Trends in Neurosciences. 39: 100–113.

20. Vashi, N., and Justice, M.J. (2019). Treating Rett syndrome: from mouse models to human therapies. Mamm Genome 30, 90–110.

21. Tiganis, T., and Tonks, N.K. (2025). Mechanisms, functions and therapeutic targeting of protein tyrosine phosphatases. Nat Rev Mol Cell Biol. 10.1038/s41580-025-00882-9.

22. Fukuhara, S., Nakajima, H., Sugimoto, S., Kodo, K., Shigehara, K., Morimoto, H., Tsuma, Y., Moroto, M., Mori, J., Kosaka, K., et al. (2019). High-fat diet accelerates extreme obesity with hyperphagia in female heterozygous Mecp2-null mice. Plos One 14, e0210184.

23. Frayre, P., Ponce-Rubio, K., Frayre, J., Medrano, J., and Na, E.S. (2024). POMC-specific knockdown of MeCP2 leads to adverse phenotypes in mice chronically exposed to high fat diet. Behav Brain Res 461, 114863.

24. Kyle, S.M., Vashi, N., and Justice, M.J. (2018) Rett syndrome: a neurological disorder with metabolic components. Open Biology. 8: 170216

25. Samaco, R.C., McGraw, C.M., Ward, C.S., Sun, Y., Neul, J.L., and Zoghbi, H.Y. (2013) Female Mecp2(+/-) mice display robust behavioral deficits on two different genetic backgrounds providing a framework for pre-clinical studies. Hum Mol Genet. 22, 96–109.

26. Goffin, D., and Zhou, Z.J. (2012) The neural circuit basis of Rett syndrome. Front Biol. 7, 428–435.

27. Jellinger, K., and Seitelberger, F. (1986). Neuropathology of Rett syndrome. American Journal of Medical Genetics Supplement 1: 259–288.

28. Bauman, M.L., Kemper, T.L., and Arin, D.M. (1995). Microscopic observations of the brain in Rett syndrome.Neuropediatrics 26: 105–108.

29. Armstrong, D.D. (2005). Neuropathology of Rett syndrome. Journal of Child Neurology 20: 747–753.

30. Guy, J., Hendrich, B., Holmes, M., Martin, J.E., and Bird, A. (2001). A mouse Mecp2-null mutation causes neurological symptoms that mimic Rett syndrome. Nature Genetics 27: 322–326.

31. Chen, R.Z., Akbarian, S., Tudor, M., and Jaenisch, R. (2001). Deficiency of methyl-CpG binding protein-2 in CNS neurons results in a Rett-like phenotype in mice. Nature Genetics 27: 327–331.

32. Krishnan, N., Koveal, D., Miller, D.H., Xue, B., Akshinthala, S.D., Kragelj, J., Jensen, M.R., Gauss, C.M., Page, R., Blackledge, M., et al. (2014). Targeting the disordered C terminus of PTP1B with an allosteric inhibitor. Nat Chem Biol 10, 558–566.

33. Krishnan, N., Konidaris, K.F., Gasser, G., and Tonks, N.K. (2018). A potent, selective, and orally bioavailable inhibitor of the protein-tyrosine phosphatase PTP1B improves insulin and leptin signaling in animal models. J Biol Chem 293, 1517–1525.

34. Kerr, A.M., Armstrong, D.D., Prescott, R.J., Doyle, D., and Kearney, D.L. (1997). Rett syndrome: analysis of deaths in the British survey. European Child & Adolescent Psychiatry 6(Suppl 1): 71–74.

35. Tarquinio, D.C., Hou, W., Neul, J.L., et al. (2015). The changing face of survival in Rett syndrome and MECP2-related disorders. Pediatric Neurology 53: 402–411.

36. Julu, P.O.O., Kerr, A.M., Hansen, S., Apartopoulos, F., Jamal, G.A., and Engerström, I.W. (1997). Immature medullary cardiorespiratory neurones in Rett syndrome. European Child & Adolescent Psychiatry 6(Suppl 1): 47–54.

37. Weese-Mayer, D.E., Lieske, S.P., Boothby, C.M., et al. (2006). Autonomic dysregulation in young girls with Rett Syndrome during nighttime in-home recordings. Pediatric Pulmonology 41: 1045–1060.

38. Singh, J., Lanzarini, E., and Santosh, P. (2020) Autonomic dysfunction and sudden death in patients with Rett syndrome: a systematic review. J Psychiatry Neurosci. 45, 150–181.

39. Sekul, E.A., Moak, J.P., Schultz, R.J., Glaze, D.G., Dunn, J.K., and Percy, A.K. (1994). Electrocardiographic findings in Rett syndrome: an explanation for sudden death? Journal of Pediatrics 125: 80–82.

40. McCauley, M.D., Wang, T., Mike, E., Herrera, J., Beavers, D.L., Huang, T.W., Ward, C.S., Skinner, S., Percy, A.K., Glaze, D.G., Wehrens, X.H.T., and Neul, J.L. (2011). Pathogenesis of lethal cardiac arrhythmias in Mecp2 mutant mice: implication for therapy in Rett syndrome. Science Translational Medicine 3: 113ra125.

41. Katz DM, Dutschmann M, Ramirez JM, Hilaire G. (2009) Breathing disorders in Rett syndrome: progressive neurochemical dysfunction in the respiratory network. Respiratory Physiology & Neurobiology. 168: 101–108.

42. Ward CS, Arvide EM, Huang TW, Yoo J, Noebels JL, Neul JL. Respiratory dysfunction in Rett syndrome: understanding epigenetic regulation of the respiratory network. Respiratory Physiology & Neurobiology. 2011;179(1):55–63.

43. Ramirez, J.M., Karlen-Amarante, M., Wang, J.J., Huff, A., and Burgraff, N. (2022) Breathing disturbances in Rett syndrome. Handb Clin Neurol. 189, 139–151.

44. Gooden BA. Mechanism of the human diving response. Integrative Physiological and Behavioral Science. 1994;29(1):6–16.

45. Schagatay E, Andersson J, Nielsen B. Human diving response and breath-hold duration in apnea divers and controls. Undersea & Hyperbaric Medicine. 2001;28(3):125–134.

46. Johnson, C.M., Cui, N., Zhong, W., Oginsky, M.F., and Jiang, C. (2015) Breathing abnormalities in a female mouse model of Rett syndrome. J Physiol Sci. 65, 451–9.

47. Cacciatore, I., Cornacchia, C., Baldassarre, L., Fornasari, E., Mollica, A., Stefanucci, A., and Pinnen, F. (2012). GPE and GPE analogues as promising neuroprotective agents. Mini Rev Med Chem 12, 13–23.

48. Parent, H., Ferranti, A., and Niswender, C. (2023). Trofinetide: a pioneering treatment for Rett syndrome. Trends Pharmacol Sci 44, 740–741.

49. Hudu, S.A., Elmigdadi, F., Qtaitat, A.A., Almehmadi, M., Alsaiari, A.A., Allahyani, M., Aljuaid, A., Salih, M., Alghamdi, A., Alrofaidi, M.A., et al. (2023). Trofinetide for Rett Syndrome: Highlights on the Development and Related Inventions of the First USFDA-Approved Treatment for Rare Pediatric Unmet Medical Need. J Clin Med 12. 10.3390/jcm12155114.

50. Kennedy, M., Glass, L., Glaze, D.G., Kaminsky, S., Percy, A.K., Neul, J.L., Jones, N.E., Tropea, D., Horrigan, J.P., Nues, P., et al. (2023). Development of trofinetide for the treatment of Rett syndrome: from bench to bedside. Front Pharmacol 14, 1341746.

51. Neul, J.L., Percy, A.K., Benke, T.A., Berry-Kravis, E.M., Glaze, D.G., Peters, S.U., Jones, N.E., and Youakim, J.M. (2022). Design and outcome measures of LAVENDER, a phase 3 study of trofinetide for Rett syndrome. Contemp Clin Trials 114, 106704.

52. Neul, J.L., Percy, A.K., Benke, T.A., Berry-Kravis, E.M., Glaze, D.G., Marsh, E.D., Lin, T., Stankovic, S., Bishop, K.M., and Youakim, J.M. (2023). Trofinetide for the treatment of Rett syndrome: a randomized phase 3 study. Nat Med 29, 1468–1475.

53. Percy, A.K., Neul, J.L., Benke, T.A., Berry-Kravis, E.M., Glaze, D.G., Marsh, E.D., Barrett, A.M., An, D., Bishop, K.M., and Youakim, J.M. (2024). Trofinetide for the treatment of Rett syndrome: Long-term safety and efficacy results of the 32-month, open-label LILAC-2 study. Med 5, 1275–1281.

54. Percy, A.K., Ryther, R., Marsh, E.D., Neul, J.L., Benke, T.A., Berry-Kravis, E.M., Feyma, T., Lieberman, D.N., Ananth, A.L., Fu, C., et al. (2025). Results from the phase 2/3 DAFFODIL study of trofinetide in girls aged 2-4 years with Rett syndrome. Med 6, 100608.

55. Sara, V.R., Carlsson-Skwirut, C., Bergman, T., Jornvall, H., Roberts, P.J., Crawford, M., Hakansson, L.N., Civalero, I., and Nordberg, A. (1989). Identification of Gly-Pro-Glu (GPE), the aminoterminal tripeptide of insulin-like growth factor 1 which is truncated in brain, as a novel neuroactive peptide. Biochem Biophys Res Commun 165, 766–771.

56. Saura, J., Curatolo, L., Williams, C.E., Gatti, S., Benatti, L., Peeters, C., Guan, J., Dragunow, M., Post, C., Faull, R.L., et al. (1999). Neuroprotective effects of Gly-Pro-Glu, the N-terminal tripeptide of IGF-1, in the hippocampus in vitro. Neuroreport 10, 161–164.

57. Sizonenko, S.V., Sirimanne, E.S., Williams, C.E., and Gluckman, P.D. (2001). Neuroprotective effects of the N-terminal tripeptide of IGF-1, glycine-proline-glutamate, in the immature rat brain after hypoxic-ischemic injury. Brain Res 922, 42–50.

58. Bickerdike, M.J., Thomas, G.B., Batchelor, D.C., Sirimanne, E.S., Leong, W., Lin, H., Sieg, F., Wen, J., Brimble, M.A., Harris, P.W., and Gluckman, P.D. (2009). NNZ-2566: a Gly-Pro-Glu analogue with neuroprotective efficacy in a rat model of acute focal stroke. J Neurol Sci 278, 85–90.

59. Li, J., Choi, E., Yu, H.T., and Bai, X.C. (2019). Structural basis of the activation of type 1 insulin-like growth factor receptor. Nature Communications 10. 4567 10.1038/s41467-019-12564-0.

60. Achilly, N.P., Wang, W., and Zoghbi, H.Y. (2021) Presymptomatic training mitigates functional deficits in a mouse model of Rett syndrome. Nature. 592, 596–600. (Erratum in: Nature. 2026 May 11. doi: 10.1038/s41586-026-10578-5).

