## Supplementary figures and images for "Inhibition of protein tyrosine phosphatase PTP1B function ameliorates pathophysiological deficits in Rett Syndrome"

### Supplementary Figures 1-3

#### Supplementary Figure 1

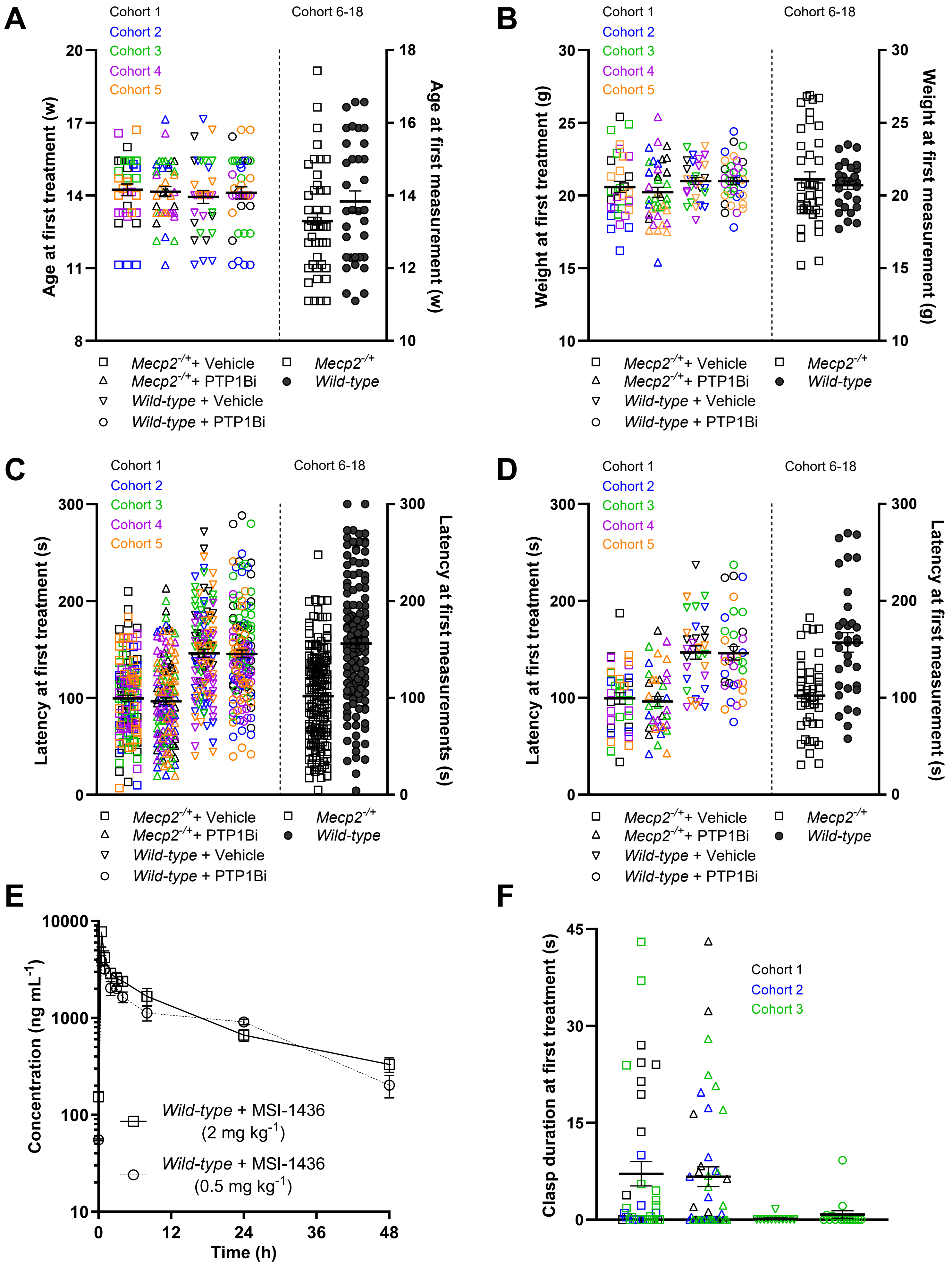

#### Supplementary Figure 2

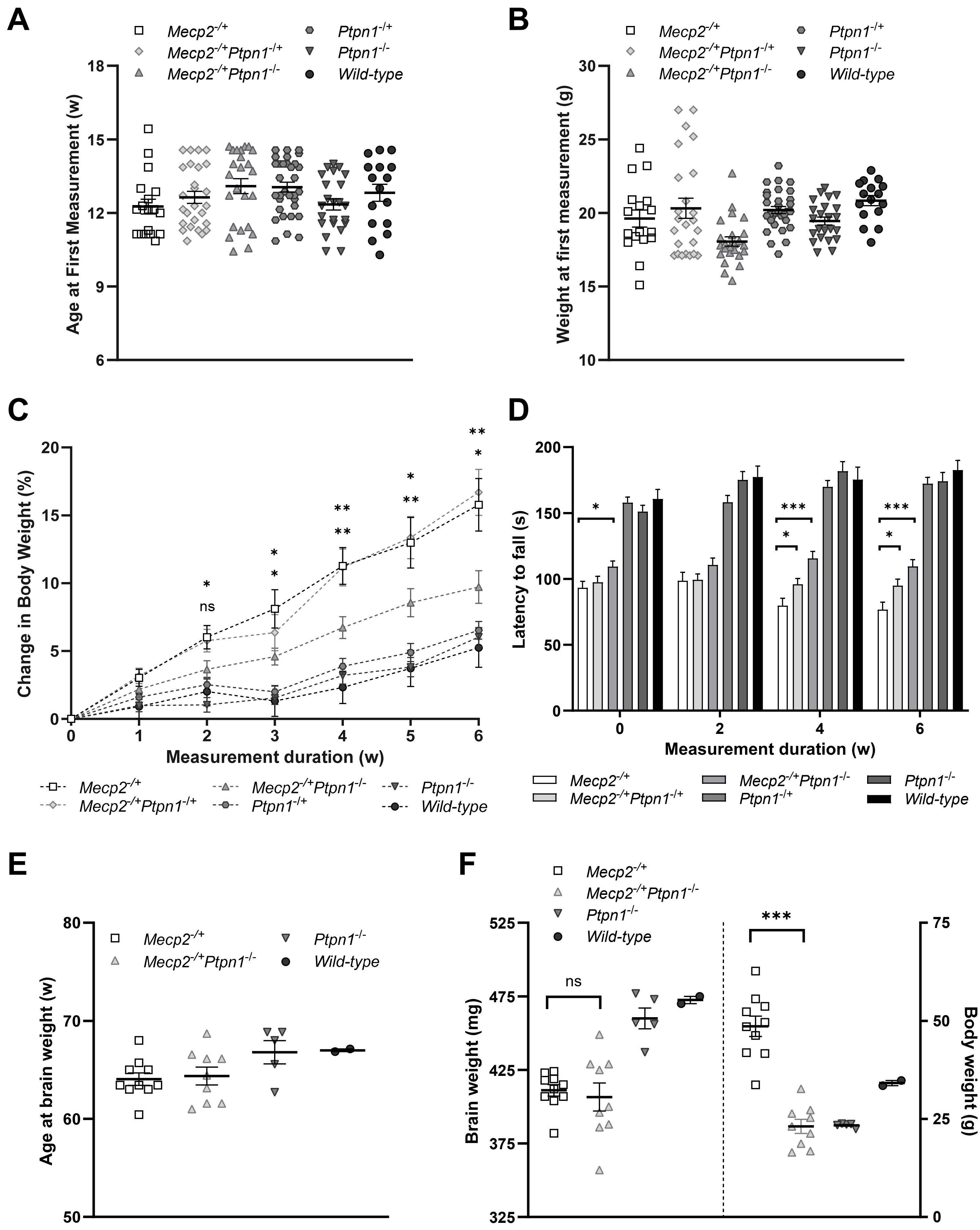

#### Supplementary Figure 3

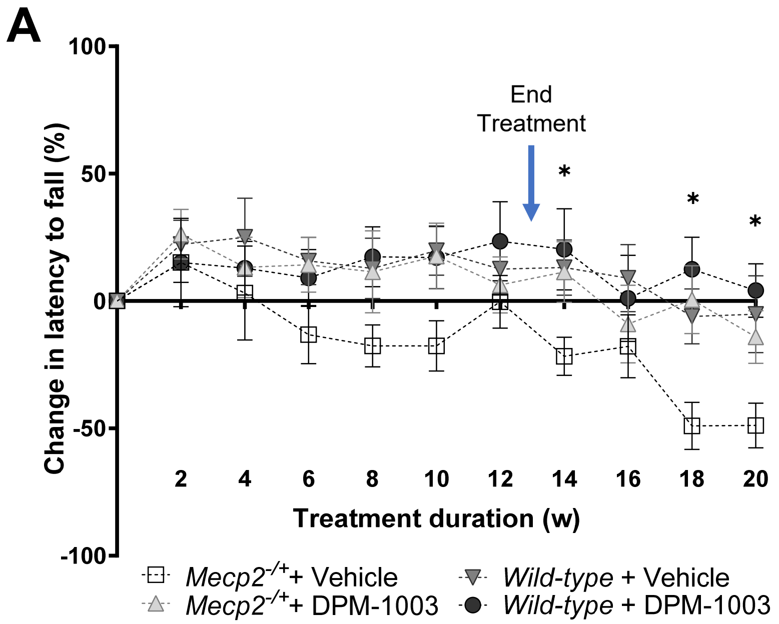
